# The soft explosive model of placental mammal evolution

**DOI:** 10.1101/251520

**Authors:** Matthew J Phillips, Carmelo Fruciano

**Affiliations:** School of Earth, Environmental and Biological Sciences, Queensland University of Technology, Brisbane, Australia

**Keywords:** Cretaceous-Paleogene boundary, fossil calibration, life history, molecular dating, Placentalia

## Abstract

Recent molecular dating estimates for placental mammals echo fossil inferences for an explosive interordinal diversification, but typically place this event some 10-20 million years earlier than the Paleocene fossils, among apparently more “primitive” mammal faunas. However, current models of molecular evolution do not adequately account for parallel rate changes, and result in dramatic divergence underestimates for large, long-lived mammals such as whales and hominids. Calibrating among these taxa shifts the rate model errors deeper in the tree, inflating interordinal divergence estimates. We employ simulations based on empirical rate variation, which show that this “error-shift inflation” can explain previous molecular dating overestimates relative to fossil inferences. Molecular dating accuracy is substantially improved in the simulations by focusing on calibrations for taxa that retain plesiomorphic life-history characteristics. Applying this strategy to the empirical data favours the soft explosive model of placental evolution, in line with traditional palaeontological interpretations – a few Cretaceous placental lineages give rise to a rapid interordinal diversification following the 66 Ma Cretaceous-Paleogene boundary mass extinction. Our soft explosive model for the diversification of placental mammals brings into agreement previously incongruous molecular, fossil, and ancestral life history estimates, and closely aligns with a growing consensus for a similar model for bird evolution. We show that recent criticism of the soft explosive model relies on ignoring both experimental controls and statistical confidence, as well as misrepresentation, and inconsistent interpretations of morphological phylogeny. More generally, we suggest that the evolutionary properties of adaptive radiations may leave current molecular dating methods susceptible to overestimating the timing of major diversification events.

## Introduction

Molecular and palaeontological analyses of placental mammals both identify an interordinal diversification spike, in which the stem lineages of nearly all 18 modern orders (e.g. primates, rodents) originated over a period of just a few million years (Ma). However, most molecular dating estimates (e.g. [1-3]) for this diversification are 10-20 Ma older than observed in the fossil record [4, 5]. The extraordinary fossil record surge for eutherians (crown placentals and their extinct stem relatives) follows the 66 Ma Cretaceous-Paleogene boundary (KPg) mass extinction event (Fig. 1). This fossil record diversification also manifests as a taxonomic phase change, with eutherians as a percentage of new mammal species appearances increasing from an average of 27% during the Campanian and Maastrichtian, to 84% during the Paleocene.

**Fig. 1.**
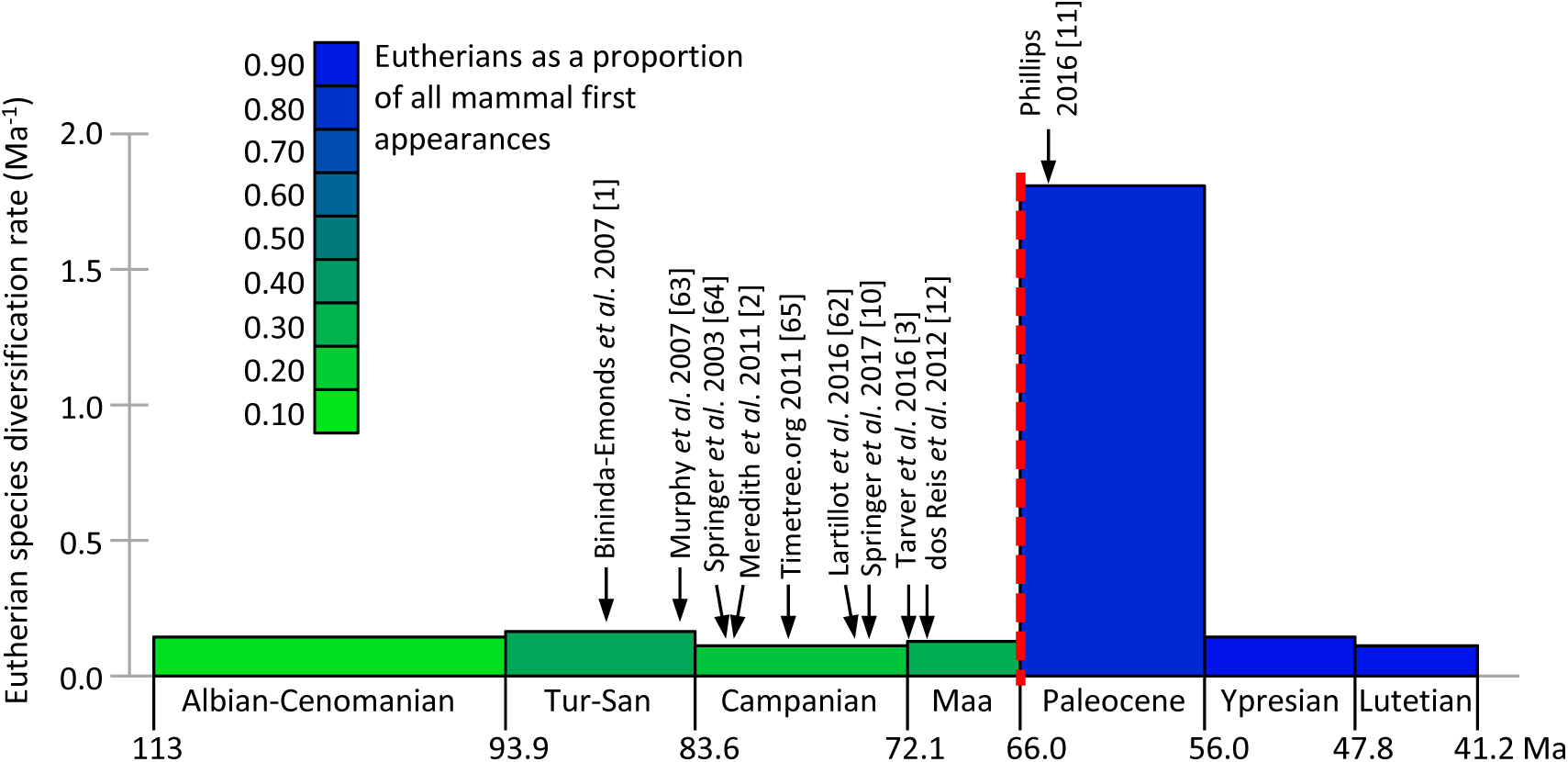
Eutherian fossil record species diversification rate. Data points are calculated as new appearances/Ma (scaled by species richness in the previous time bin, a proxy for starting species richness). Barremian-Aptian provides the previous time bin for Albian-Cenomanian. The green-blue colour shading indicates the proportion of eutherians among mammal first appearances for each time bin. Referenced arrows indicate molecular dating estimates for the temporal midpoint of the placental interordinal diversification, including for Laurasiatheria, Euarchantoglires and Afrotheria (See Supplemental file 3: Table S3). The KPg boundary is shown as red dashes. Tur-San, Turonian to Santonian; Maa, Maastrichtian.

If the older molecular dates for the interordinal diversification are instead correct, then this most profound event in placental history leaves no discernible trace in the fossil record (Fig. 1). This is especially perplexing, because ancestral area reconstruction [6] places this proposed ∼75-85 Ma molecular radiation (including stem members of all 11 Northern Hemisphere orders) right alongside the best Late Cretaceous fossil faunas in Eurasia and North America. It is similarly incongruous that during the diversity surge in the placental fossil record following the KPg mass extinction [7, 8] those same molecular timetrees instead imply stable or even declining diversification [2, 9]. Springer *et al*.’s [10] new tree does place several additional divergences close to the KPg relative to [2], but this appears to be an artefact of adding maximum bounds at the KPg for these clades to bump up against.

Phillips [11] recently presented evidence for two methodological contributors to molecular dates overestimating early divergences among placentals: (1) Molecular clocks over-smooth parallel decelerations in evolutionary rates among large, long-lived mammals. This results in several-fold divergence underestimates in groups such as whales and seacows, for which calibration to correct these clade ages simply transfers the underlying rate error stemwards, and inflates divergence estimates deeper in the tree. (2) Such “error-shift inflation” is further facilitated by asymmetry in calibration priors between minimum bounds that are highly speculative, and maximum bounds that are too conservative to buffer against rate misspecification or erroneous minimum bounds at other nodes.

Phillips [11] sought to ameliorate error-shift inflation in two steps. The first reduced the impact of oversmoothed, parallel rate decelerations on dates deeper in the tree, by employing dos Reis *et al.*’s [12] calibration scheme – which includes fewer constraints among large, long-lived taxa than does Meredith *et al.*’s [2] scheme. The second step reduced asymmetry in fossil calibration priors, by revising overly conservative maximum bounds in line with best practices [13], so as to enhance the capacity of the calibration scheme to buffer against rate errors. The revised calibration scheme was then used to reanalyse Meredith *et al.*’s [2] 26-locus dataset for 169 taxa, and resulted in molecular dates that closely matched long-held fossil record expectations [14-16]. We refer to this as the “soft explosive” model of placental evolution; a few Cretaceous placental lineages seed the massive interordinal diversification spike that follows the KPg extinction event.

Although the soft explosive model brings agreement between molecular and fossil inference of placental evolution, it has recently been criticized by Springer *et al*. [10] on two grounds. The first criticism is that Phillips [11] erroneously dragged divergences younger by “eliminating calibrations in large-bodied/long lifespan clades” without deleting those taxa. This claim is false. Phillips [11] maintained each of dos Reis *et al.*’s [12] calibrations that were placed in large-bodied/long lifespan clades. Springer *et al.*’s [10] argument was also based on an analysis in which they deleted large, long-lived taxa, and found that most supraordinal divergences increased by 8-10 Ma relative to Phillips [11]. However, Springer *et al.* [10] failed to control for calibration, and it is not their taxon deletion, but their inclusion of poorly supported calibrations that drives the divergence estimates older (as discussed below; also see Supplemental file 1). Indeed, when we repeat their taxon deletion, but maintain the original calibration scheme of Phillips [11], the divergence estimates again support the soft explosive model (dR32 analysis, Table 1C).

**Table 1.**
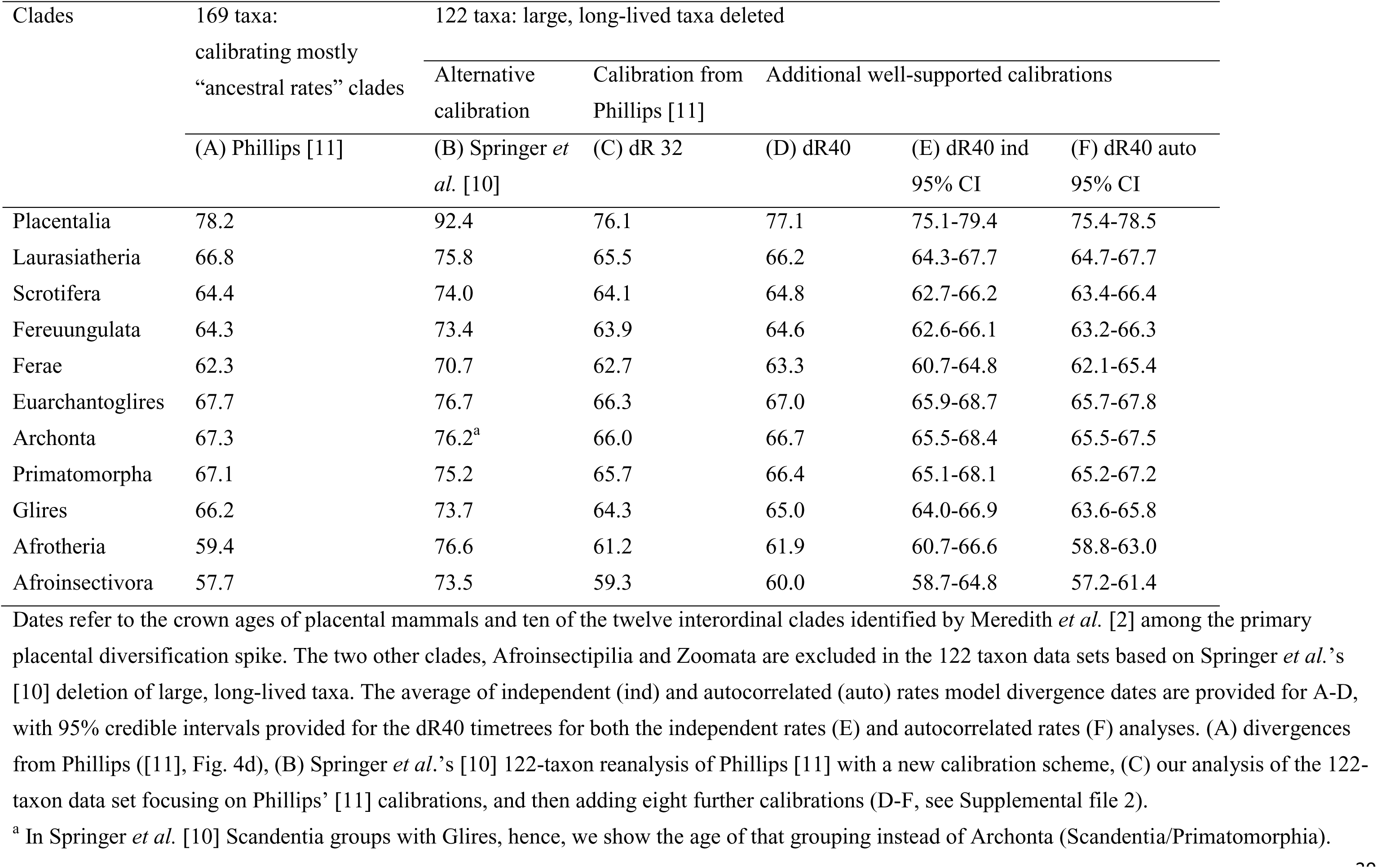
Mean MCMCtree posterior estimates (Ma). Clades 169 taxa:calibrating mostly

Springer *et al.*’s [10] second criticism of the soft explosive model was that many divergences post-date the earliest fossil evidence for the clade, thus implying the existence of fossil “zombie” lineages. We will show that this criticism is based on misrepresentation, false precision in their interpretation of molecular dates, and overconfidence in poorly resolved fossil relationships.

One point of agreement in the debate over the timescale of placental evolution is that calibrating among large, long-lived taxa results in older age estimates for the root and interordinal divergences [2, 10, 11]. Nevertheless, attempting to understand and quantify the contribution from error-shift inflation may be confounded by variation in calibration precision (how closely fossil calibrations match true divergences) – which may differ between the deleted large, long-lived calibrations and the remaining calibrations. Here we use simulations based on empirical estimates of molecular rate variation among placentals to control for calibration accuracy and precision, and to better understand the influence of error-shift inflation.

Considered together, our simulation study and new molecular dates based on revised fossil calibrations for the datasets of Meredith *et al.* [2] and Liu *et al.* [20] provide strong support for the soft explosive model of placental diversification. Moreover, previous, older molecular dates are explained as artefacts of errors in both calibration and modelling rate variation across the tree. In turn, the younger KPg diversification allows us to revise Romiguier *et al.*’s [18] surprising molecular inference of early placental life history traits. More generally, there is a wider pattern of conflict between molecular dates and fossil evidence for the timing of major diversifications, such as for birds [19, 20], flowering plants [21-23] and the Cambrian explosion [24]. We discuss the possibility that major adaptive radiations could be particularly susceptible to error-shift inflation, resulting in molecular divergence overestimates.

## Results and Discussion

### Simulated rate deceleration among large, long-lived taxa mimics observed molecular dating errors

To control for calibration and isolate the behaviour of error-shift inflation, we simulated molecular data on a phylogeny of given age (Fig. 2A) that is simplified from the proposed mammalian timetree of Phillips ([11], Fig. 5). We simulate “placentals” originating at 80 Ma, then splitting into two 66 Ma superorders (e.g. “Laurasiatheria” and “Afrotheria”), which each give rise to two 33 Ma calibrated clades. In the first set of simulations the branch rates are randomly drawn from a lognormal distribution modelled on inferred rates from Phillips [11] for small to mid-sized mammals (<30kg adult body mass, <40 years maximum longevity). Relaxed molecular clock dating in MCMCtree [25, 26] accurately reconstructs all node heights under this simple distribution of rates across the tree (Fig. 2C-E, light grey bars). However, when we simulate a parallel rate deceleration reminiscent of whales or seacows for just one of the 33 Ma calibrated nodes in each superorder, the MCMCtree reconstructions reveal extreme error-shift inflation. Average estimates for the 66 Ma superorders were inflated to 80.5 Ma, and the 80 Ma placental root was inflated to 107.9 Ma. In each case the 95% CIs are fully older than the simulated dates. These inflated divergences closely mimic recent molecular dates for placental mammals.

**Fig. 2.**
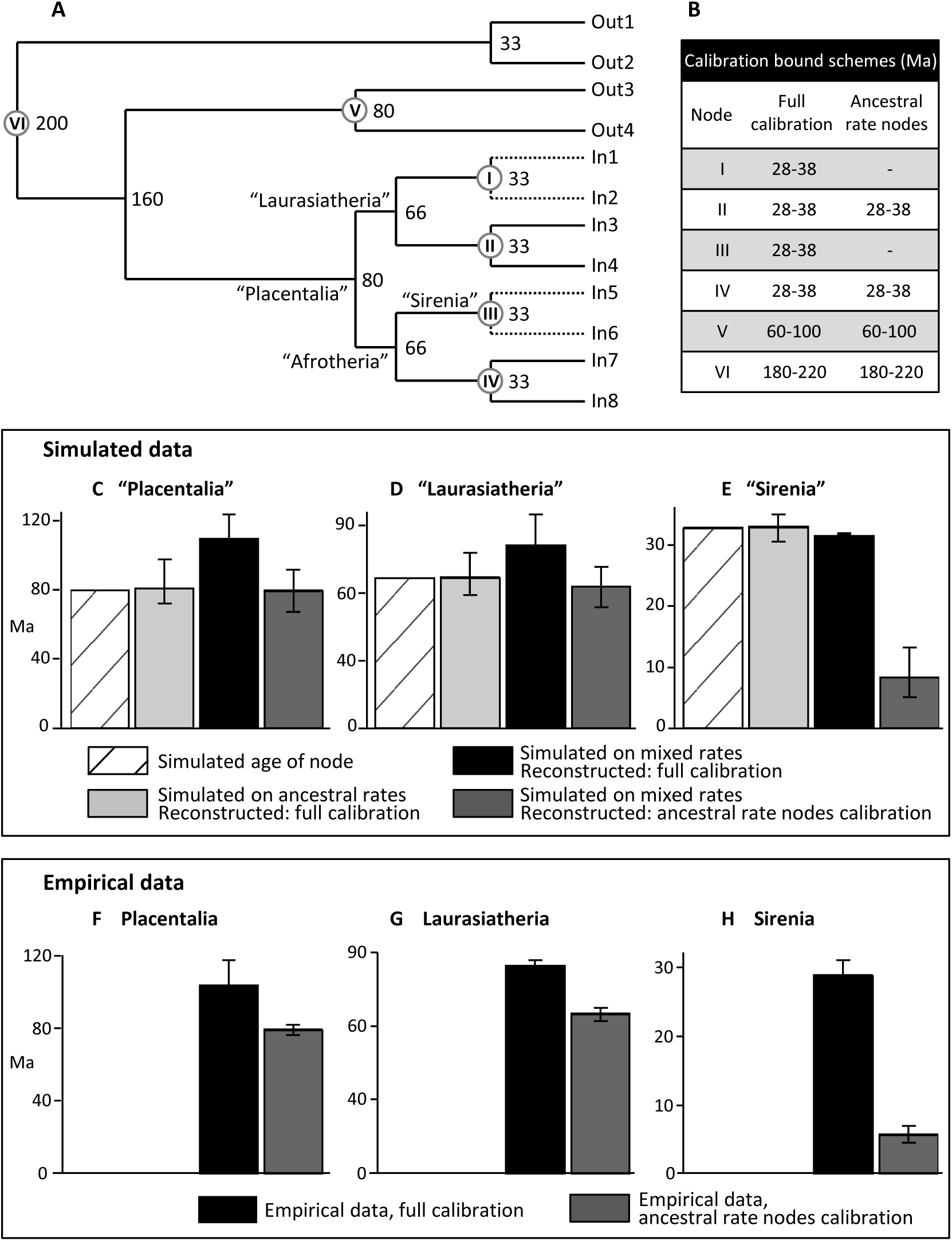
Simulating error-shift inflation of deep placental divergences, and amelioration by excluding calibrations among large, long-lived clades. A. Dated tree on which 20,000 bp DNA sequences were simulated under two rate schemes, “ancestral rates” with all branch rates drawn from a single lognormal distribution (see Materials and Methods) and “mixed rates” with the rate drawn from that same distribution, except divided by 5 for the daughter lineages of nodes I and III. B. Soft bound calibrations on nodes I-VI under alternative MCMCtree reconstructions. Date estimates and 95% CIs for simulated clades (C) “Placentalia” (D) “Laurasiatheria” and (E) “Sirenia” are shown for the “ancestral rates” simulation (light grey), and for the “mixed rates” simulation with either full calibration (black) or calibration only on ancestral rates clades (dark grey). Corresponding date estimates from Phillips ([11], Fig. 4c,d) on the empirical data are shown for (F) Placentalia, (G) Laurasiatheria and (H) Sirenia, with full calibration (black) and largely focusing on ancestral rates clades (dark grey).

Our simulations also reproduce the empirical pattern of extreme dating underestimation for large, long-lived clades for when they are not calibrated. In particular, the low rate clades simulated as 33 Ma are reconstructed by MCMCtree with a mean age of 8.5 Ma, almost as extreme as the empirical pattern for seacow origins falling from ∼28 Ma to 5.7 Ma when uncalibrated (Fig. 2E,H). Importantly though, excluding calibrations among the low rate (large, long-lived) clades allows accurate inference of divergence dates deeper in the tree, returning reconstructions close to the simulated ages (Fig. 2C,D “mixed rates ancestral calibrations” – dark grey bars). It is remarkable how closely the pattern of uncalibrating large, long-lived taxa to overcome the simulated error-shift inflation (Fig. 2C-E) mirrors the empirical pattern for placental mammals (Fig. 2F-H). Thus, our simulations, which are based on empirical rate variation, show that error-shift inflation associated with parallel rate deceleration among large, long-lived placentals can explain the proposed overestimation of interordinal divergences among molecular dating analyses.

### Conjuring up “zombie” lineages

Lane *et al.* [27] coined the term “zombie lineage” for the extension of a taxon’s survival beyond their last fossil appearance. Springer *et al.* [10] re-purposed the term for molecular divergences that are younger than minimum ages implied by fossil records, and claim that Phillips’ [11] “preferred timetree” includes 61 (of 136) internal placental nodes that are younger than first fossil appearances, thus resulting in “zombie” lineages. Springer *et al.*’s [10] claim is based on a series of misrepresentations, which are best appreciated by first understanding how Phillips’ [11] timetree was constructed. Phillips [11] recognised that calibrating large, long-lived taxa in the tree of more plesiomorphic mammals erroneously inflates interordinal divergences (also shown here with simulations, Fig. 2C,D “full calibration” – black bars), whereas not calibrating among these taxa underestimates their own within-family divergences (Fig.2E “mixed rates ancestral calibrations” – dark grey bar). Phillips [11] addressed this challenge in two steps. The first step inferred divergences with dos Reis *et al.*’s [12] calibrations, most of which are set among taxa with plesiomorphic life-history (tree 1, Fig. 4d in [11]). The final timetree (tree 2, Fig. 5 in [11]) was then inferred with more calibrations added among large, long-lived taxa, but with maximum bounds on several superordinal clades based on broad agreement between tree 1 and fossil records for major diversification following the KPg (also noting that multi-lineage diversifications should provide more robust markers in the fossil record than individual nodes).

Springer *et al.*’s [10] misrepresentation begins by overlooking Phillips’ [11] discussion of uncalibrated divergences among large, long-lived taxa being underestimated in tree 1, and falsely claiming tree 1 to be Phillips’ [11] “preferred tree”. They then ignore the final timetree with those taxa calibrated (tree 2), which Phillips [11] used for final inference of molecular rates, and instead, Springer *et al*. [10] set up the tree 1 dates as a straw man for comparison with fossil dates.

A careful examination of the (actually) 62 nodes that Springer *et al.* [10] tabled as postdating proposed fossil dates reveals that 40 involve clades of large/long-lived taxa. These underestimates follow directly from the aims for tree 1, which were to reveal the extent of date underestimation among large, long-lived clades and to isolate the interordinal nodes from error-shift inflation that would result from those large, long-lived clades being calibrated (as our simulation study confirms, Fig. 2). Springer *et al.* [10] perhaps agree, and deleted all 40 of those large, long-lived clades for their analysis.

The more concerning claim that Phillips [11] underestimated the age of 22 clades that retain apparently more plesiomorphic life history traits is illusory, created from false precision. Springer *et al.* [10] exaggerate disagreement here by treating those molecular dates as errorless, and by ignoring uncertainty in the phylogenetic attribution of reference fossils for minimum bounds. If instead we undertake the usual practices of basing fossil minimum dates on phylogenetically well-supported fossils, and considering Bayesian molecular divergences as 95% credible intervals (CIs), then the discrepancy vanishes for 20 of the 22 clades (Supplemental file 1). Moreover, Phillips’ [11] final molecular dates (tree 2) place the 95% CIs for the two remaining clades, Musteloidea (28.5-30.7 Ma) and Feliformia (30.3-35.3 Ma) entirely older than their respectively proposed fossil minima of 24.8 Ma and 28.1 Ma. The tree 2 analysis was primarily designed to test rate variation hypotheses, and still retains some dubious fossil calibrations from Meredith *et al*. [2]. But it is notable that our primary dating estimates in this study are also consistent with both of these proposed fossil minima (Supplemental file 3, 122-taxon dR40 trees). Thus, Springer *et al.*’s [10] claim of “zombie” lineages among smaller, shorter lived taxa is unfounded.

### Dating the evolution of placental mammals

We have updated our calibration set to allow for eight additional well-supported calibrations (Supplemental file 2) that were not employed by dos Reis *et al.* [12], but include several that Springer *et al.’s* [10] list of “zombie” lineages implied would increase our divergence estimates. This lifts the number of calibration priors to 40 for the same 122 taxa with apparently plesiomorphic life histories that were employed by Springer *et al.* [10]. The resulting MCMCtree timetree (dR40, see Table 1D) provides very similar divergence estimates to our previous calibration schemes (Table 1A,C). The most profound diversification in placental mammal history again falls across or closely follows the KPg boundary (see Fig. 3A), including for the origin and basal radiation of all three major superorders (Laurasiatheria, Afrotheria and Euarchontoglires).

**Fig. 3.**
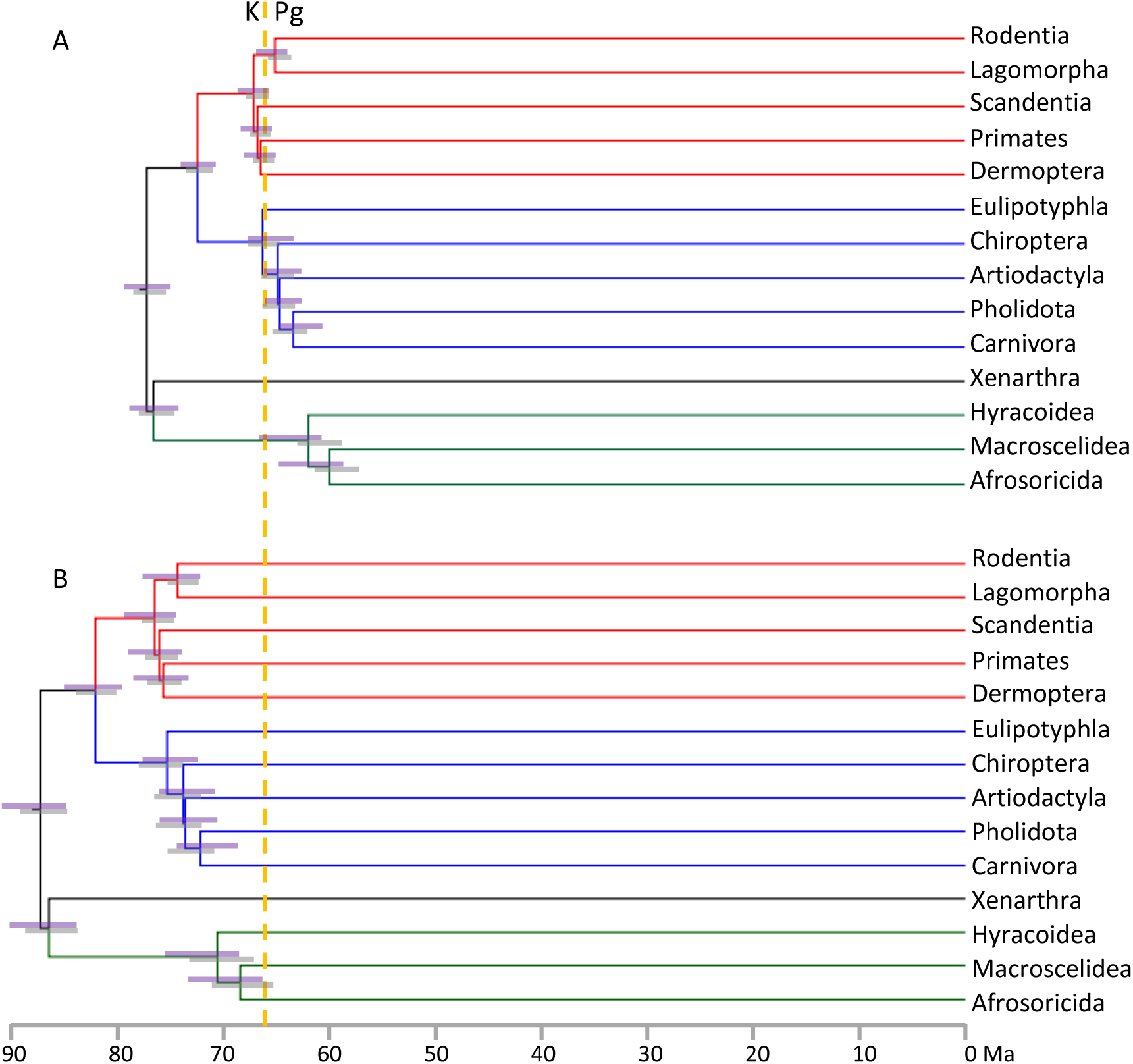
Placental mammal ordinal-level timetrees on the 122-taxon dataset for which large, long-lived taxa are excluded. Node heights are averaged over MCMCtree independent and autocorrelated rates analyses, with 95% CIs shown for analyses under independent rates (purple bars) and autocorrelated rates (grey bars). A. using our dR40 calibration set. B. Adding additional poorly-vetted calibrations for Lorisiformes, Lagomorpha and Emballonuroidea, and with maximum bounds for Primates, Rodentia and Chiroptera increased to the KPg, following Springer *et al.* [10]. Adding the additional calibrations inflates the midpoint for the primary placental interordinal diversification from 64.6 Ma to 73.7 Ma. This is largely attributable to the primate, rodent and lagomorph calibrations. Adding the emballonuroid calibration alone to our dR40 analysis only shifted the midpoint from 64.6 Ma to 65.7 Ma.

To isolate the source of the differences between our dates and Springer *et al.*’s [10] dates we identified poorly-vetted reference fossils that they used to define eight placental calibration minima that are older than our dR40 molecular estimates (Table 2). In most cases the temporal difference is so minor (0.1-2.3 Ma) as to have little impact deeper in the tree. However, three of Springer *et al.*’s [10] fossil minima are strikingly older than our molecular estimates, and reveal stark inconsistency in how these authors treat morphological phylogenetic evidence. For example, Springer *et al.* [10, 28, 29] express valid cautions, and are highly critical of morphological phylogeny, even for well-sampled modern or Mesozoic eutherians that are analysed within objective, statistical frameworks. Yet, when it comes to calibration, Springer *et al.* [10] accept reference fossils based on highly fragmentary material, unverified by any formal phylogenetic analyses (matrix-based or otherwise) or that are contradicted by such analyses (see Supplemental file 1) – then employ these fossils as minimum bounds with 97.5% or indeed, preferentially with 100% prior probability.

**Table 2.**
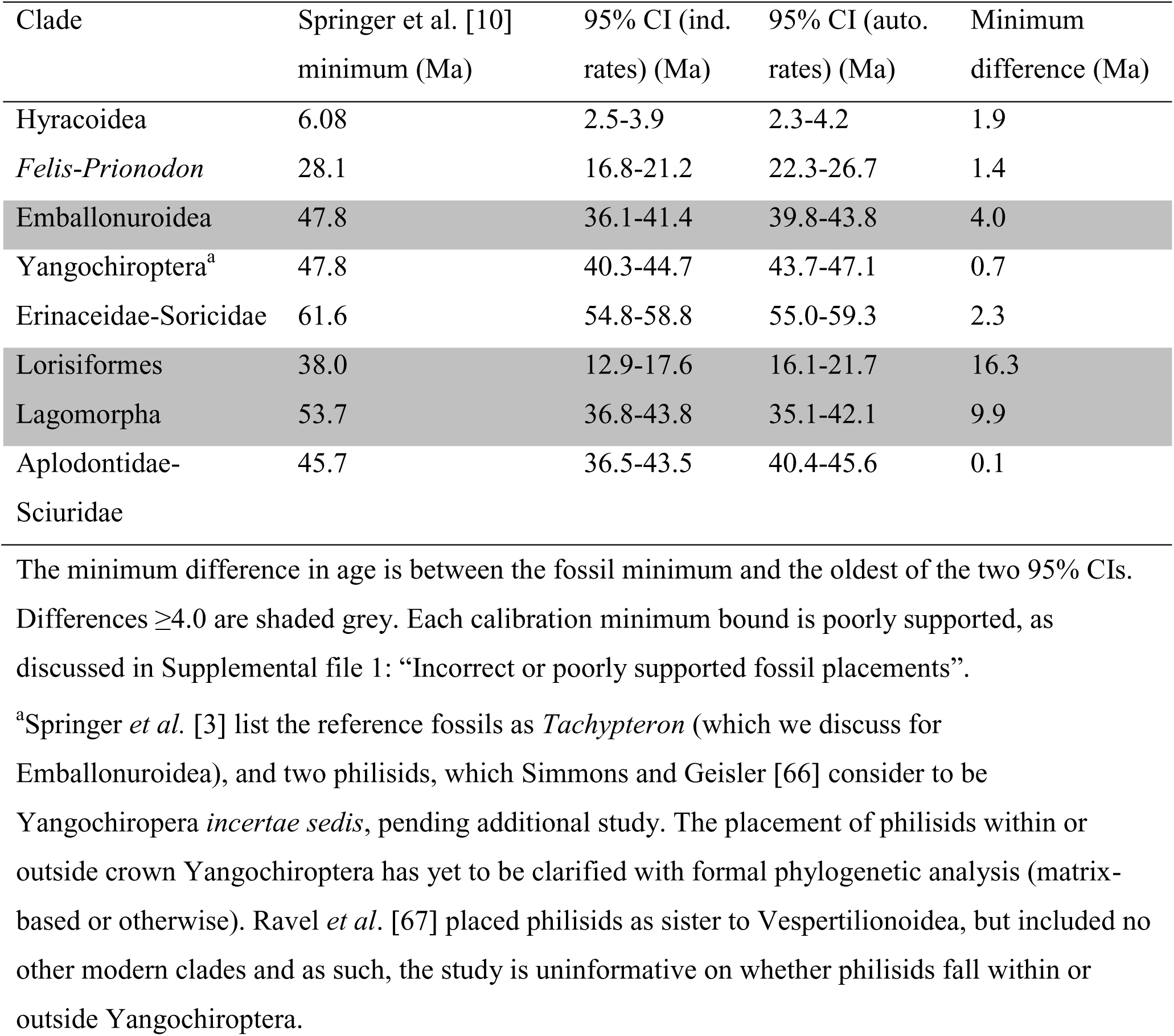
Springer *et al.*’s [10] placental calibration minima that are older than our 95% CIs for both the independent and autocorrelated rates models, using our dR40 calibration set.

The three poorly-vetted fossils that substantially conflict with our molecular dates are:

1. The ∼38 Ma *Saharagalago* (calibrating Lorisiformes) is known from just two molars. Phillips ([11], Fig. 2) showed this calibration to be an extreme outlier for apparent dating error (or rate distortion). The most likely explanation is that *Saharagalago* (and *Karanisia* from the same locality) falls outside Lorisiformes, as two recent phylogenetic analyses found [30, 31].
2. The ∼53 Ma ‘Vastan’ ankle bones (calibrating Lagomorpha) were found by Rose *et al.* [32] to group with *Oryctolagus* to the exclusion of other rabbits and hares. If true, these ankle bones would implausibly pre-date molecular dating expectations for the *Oryctolagus-Sylvilagus* divergence by ∼5-fold [33]. However, Rose *et al.* [32] did not consider sampling error and noted that the morphological signal may be confounded by functional similarities. Zhang *et al.*’s [34] μCT scans have since shown that a key character, the calcaneal canal, is also present in stem lagomorphs. Our re-analysis of Rose *et al.* [32] excluding the calcaneal canal character places the Vastan ankle bones outside crown lagomorphs, although their affinities remain statistically unresolved (Supplemental file 1: Figure S1).
3. The ∼47 Ma bat, *Tachypteron* (calibrating Emballonuroidea) was assigned by Storch *et al*. [35] only on the basis of similarities, and within a framework that considered emballonurids as sister to rhinolophoids. These two groups are now known to fall on opposite sides of the chiropteran tree [36], and some other Eocene European bats previously assigned to Emballonuridae and Rhinolophoidea have since been placed in a new family of uncertain affinities [37]. *Tachypteron* was not considered in that study. Ravel *et al.*’s [38] cladistic analysis of *Tachypteron* and *Vespertiliavus* included only emballonuroids among crown bats. Hence, the placement of *Tachypteron* requires confirmation, especially analysis of cranial and post-cranial material.

The potential for interordinal divergences to be inflated by poorly-justified calibration minimum bounds (or by rate model errors) is exacerbated by overly conservative maximum bounds. Maximum bounds should cover the time back until relatively well sampled fossil assemblages in potential geographic regions of origin that contain no putative crown group members, but contain stem members or ecological equivalents [13, 39]. These conditions are met for Chiroptera in the Thanetian (base = 59.2 Ma) [40] and for both Primates and Rodentia in the Selandian (base = 61.6 Ma) [11]. Springer *et al.* [10] extend each of these maximum bounds by one or more additional geological stages, based on arbitrary phylogenetic bracketing from [41] or unspecified uncertainty in the case of bats, from [42].

It is now apparent that the few most problematic minimum and maximum calibration bounds listed above are the main drivers for Springer *et al.* [10] pushing the primary diversification of placental mammals back into the Cretaceous. This can be shown by starting with our dR40 analysis of Springer *et al.*’s [10] 122-taxon dataset, and then substituting in their minimum bounds for Lorisiformes, Lagomorpha and Emballonuroidea, and their maximum bounds for basal rodents, primates and bats. As a result, the midpoint of the primary placental interordinal diversification shifts from 64.6 Ma, back to 73.7 Ma (Fig. 3), closely matching Springer *et al.*’s [10] 74.1 Ma diversification midpoint. In contrast, our basal Paleocene estimate is in phase with the primary diversification in the fossil record (Fig. 1) and with a new generation of morphological/total evidence dating [43, 44].

One recent genome-scale inference of mammal divergence [17] is remarkably sensitive to rate model assumptions, such that their results are difficult to place on Fig. 1. Even for their favoured STAR tree and gene-wise partitioning for MCMCtree, the primary interordinal diversification midpoint varies from 68.0 Ma with independent rates to 94.7 Ma with autocorrelated rates. Further substantial dating differences across partitioning schemes might also raise possible concerns about the underlying data (also see [45]), but two other issues are worth considering within our present context. Liu *et al.* [17] calibrate several very large, long-lived mammal clades, and 16 of 19 maximum bounds were defined by the presence of a stem lineage taxon (often the oldest, and with variously putative to well-agreed support). Maximum bounds should never be based on a specific fossil taxon – a practice that cannot account for sampling artefacts, and yet, when there is a good fossil record, can depend less on the age of the crown group being calibrated and more on the divergence from its sister taxon.

We employed Liu *et al*.’s [17] genomic data with our dR40 calibration scheme on the relevant nodes after deleting the large, long-lived taxa. The resulting timetrees (Supplemental file 3) provide far closer agreement between independent and autocorrelated rates models. Liu *et al.* [17] favoured the independent rates model over autocorrelated rates, based on several simulated and empirical tests. Under the independent rates model the placental diversification midpoint is 63.2 Ma, and the initial divergences within Laurasiatheria, Euarchantoglires and Afrotheria closely co-occur (all within 2.3 Ma) instead of being spread over 7.6 Ma as in Liu *et al.* [17]. Hence, with more rigorous calibration and reducing the potential for error-shift inflation, genome-scale data support the soft explosive model of placental evolution.

### Molecular rates and life history traits among early placental mammals

Focusing calibration on clades that maintain ancestral evolutionary rates (or life history rate correlates) is the critical element shared by our most accurate dates for the simulated data and our empirical estimates for a placental origin younger than 80 Ma and major diversification near the 66 Ma KPg event (Fig. 2). This finding was foreshadowed by Phillips [11] showing that molecular rates for placental, marsupial and monotreme stem lineages were reliably traced back into the Mesozoic when calibrating clades that retain inferred ancestral life history traits, whereas calibrating only among large, long-lived mammals resulted in implausibly old divergences.

Inference of life history rate correlates from fossils also predicts that early eutherians had at least moderate rates of molecular evolution. All of the thousands of eutherian fossils from the period (Albian-Campanian: 113-72.1 Ma) that covers nearly all molecular date estimates for the origin and subsequent interordinal diversification of placentals were from small animals (<250g adult body mass) [46, 47]. Lifespans of these extinct eutherians were also likely to have been relatively short, because maximum longevity among all similarly small modern, non-volant and non-fossorial placentals is less than 20 years (mean 7.2 years; 95% CI 2.7-17.9 years, AnAge Database [48]).

One molecular argument against short longevity and high molecular rates among early placentals needs to be addressed. Romiguier *et al.* [18] analysed genomic protein coding GC content at 3^rd^ positions (GC3) and found a remarkable correlation between GC3 conservation and longevity. They estimated maximum longevity of 25.7-40.9 years for early placentals, which is well beyond the range noted above for modern eutherians that are as small as their Albian-Campanian counterparts. However, Romiguier *et al.*’s [18] GC3 conservation metric is a function of time since divergence, and they assumed that crown placentals originated at 105 Ma.

Romiguier *et al.* [18] presented a time-correlated index of GC3 conservation, γ = −t/log(τ), where t is time since divergence and τ is Kendall’s correlation coefficient for GC3 conservation among genes, between species. We recalculated γ for each of Romiguier *et al.*’s [18] GC3 conservation coefficients (τ) for taxon pairs, but with divergence estimates from Phillips [11]. We confirm the strong correlation between γ and maximum longevity (R^2^=0.91; maximum longevity = 0.0683γ – 10.243). This relationship allows divergence estimates for the origin of placental mammals to be cross validated against life history inferences drawn from the fossil record. If we use our 77 Ma estimate for the placental origin, the maximum longevity estimate for early placentals falls dramatically, to 7.9-21.9 years (Supplemental file 4), and is now consistent with many modern, small placentals. Thus, the emerging picture is of placental mammals with size and longevity similar to tree shrews, inheriting the post-KPg world and rapidly diversifying into the ecospace opened up by the extinction of dinosaurs and many other land vertebrates.

### Molecular dating adaptive radiations

The KPg rate spike that Springer *et al*. [29] claim for explosive models was produced by forcing a “hard” explosive model that compresses the placental origin and >15 Ma of evolution on our “soft” explosive tree (Fig. 3A) into just 200,000 years – an extreme scenario that they dismissed (but see [49]). In contrast, the “soft” explosive model places the interordinal diversification (not the placental origin) at the KPg and molecular rate estimates for placentals remain much the same across the KPg [11]. However, parallel rate slowdowns occur in large-bodied, long-lived clades, such as whales and seacows [11], which upon calibration provide strong upwards pressure on interordinal divergences. Similar rate-shift inflation may be promoted in birds by parallel rate slowdowns, for example, among penguins and tubenoses [50].

We expect that the three key elements of error-shift inflation will often be associated with adaptive radiations. The first is that evolutionary races into novel ecospaces, which involve negotiating complex fitness landscapes, will favour species with large effective population sizes and high substitution rates [51-53], and these will typically be smaller, shorter-lived species. Much the same is predicted by theory around Cope’s rule [54, 55] for the tendency for radiations to proceed from smaller to larger body size. The second element, is that once large body size does evolve, fossil sampling improves [56] and allometry drives apomorphy [57]. These factors tend to promote tighter minimum bounds, which combined with the rate deceleration concomitant with large body size, provides the basis for error-shift inflation. The third factor that is typical for adaptive radiations is that maximum bounds are often necessarily conservative for calibrations deeper in the tree, if they rely on detecting smaller, more plesiomorphic taxa. This in turn reduces the effectiveness of these maximum bounds for buffering against error-shift inflation associated with underestimation of parallel rate deceleration among large, long-lived taxa.

In the present study our simulations based on empirical rate variation show that error-shift inflation associated with parallel rate deceleration among large, long-lived placentals can explain the proposed overestimation of interordinal divergences among molecular dating analyses. We have overcome error-shift inflation by focusing taxon sampling (or calibration) on mammals with more plesiomorphic life history rate-correlates, and by reducing asymmetrical confidence in assigning minimum and maximum calibration bounds. As a result, the most profound diversification event in placental mammal history is brought into temporal agreement between molecular dates and the fossil record (Fig. 1). A similar soft explosive model of diversification immediately following the KPg is now emerging among birds, within both Neoaves [19, 50, 58] and palaeognaths [59]. Better understanding the relationship between natural history rate-correlates and calibration strategies may be important for resolving molecular dating/fossil record controversies for other adaptive radiations, such as for the Cambrian explosion of metazoans, and for flowering plants.

## Materials and Methods

### Simulating molecular rate evolution and error-shift inflation among placental mammals

We simulated mammalian molecular data to understand whether realistic patterns of molecular rate variation, including parallel rate decelerations among large, long-lived taxa could explain interordinal divergence overestimates, when controlling for calibration. For each set of simulations we used Seq-Gen 1.3.3 [60] to generate 100 datasets of 20,000 bp sequences for a 12-taxon phylogeny (Fig. 2A) that is simplified from the proposed mammalian timetree of Phillips [11]. In addition to “monotreme” and “marsupial” outgroups, the “placental” ingroup has its crown origin at 80 Ma, with two daughter nodes at 66 Ma (mimicking superordinal divergences, such as Laurasiatheria and Afrotheria), and each splitting into two 33 Ma clades.

In the first set of simulations, which we refer to as “ancestral rates”, the branch rates are randomly drawn from a lognormal distribution (ln mean −6.523, s.d. 0.274) modelled on inferred rates for small to mid-sized mammals (<30kg adult body mass, <40 years maximum longevity) from [11], based on the 26-gene, 169-taxon dataset of Meredith *et al*. [2]. A second set of simulations that we refer to as “mixed rates” draws from the “ancestral rates” distribution for most of the tree, but mimics large, long-lived taxa for two 33 Ma clades diverging from nodes I and III in Fig. 2B. These rates are drawn from the same lognormal distribution, but scaled to 1/5, similar to whales or seacows, from Phillips [11].

Timetrees for each simulated dataset were inferred separately in MCMCtree [25, 26], using the independent rates model. Calibrations were all symmetric, with 2.5% soft bound minima and maxima equidistant from the “true” simulated age. These age bounds are shown in Fig. 2B for all calibrated nodes. The simulated datasets were analysed either with all six calibrations (full calibration) or without calibrating the two clades (I & III) that exhibit the rate deceleration (ancestral rate nodes calibration).

### Empirical data and deleting large, long-lived taxa

Mammalian timetrees were estimated from two DNA datasets based on the 26-gene (35,603 bp), 169-taxon alignment of Meredith *et al*. [2]. The first is the 122-taxon dataset, for which Springer *et al.* [10] had deleted all taxa included by Meredith *et al.* [2] that are >10 kg and/or >40 years maximum longevity. The second dataset (128 taxa) includes additional mammals up to 30 kg, to test the sensitivity of the date estimates to including medium sized mammals well outside the upper size bound of any Mesozoic eutherians, but that are not especially long lived (still < 40 years maximum longevity). In addition, we estimated timetrees from Liu *et al*.’s [17] three favoured “first quintile” 200-gene alignments, again including only the 57 taxa that are ≤10 kg and ≤40 years maximum longevity.

The fossil calibration bounds employed for each of the empirical timetree analyses are provided in Supplemental file 2: Table S2. To summarize, our initial analysis of the dataset for 122 taxa with presumed plesiomorphic life histories employs the calibration scheme of Phillips [11], except for calibrations on nodes deleted by Springer *et al.* [10]. Most placental mammal calibrations were originally based on [12]. Next we added eight additional, well-supported calibrations, including from among those that Springer *et al.* [10] implied would increase our divergence estimates, lifting the number of calibrations to 40. A further three calibrations were added upon the inclusion of additional taxa up to 30kg for the 128-taxon dataset. Twenty-four of our favoured calibrations were compatible with the taxon sampling for the alignments derived from Liu *et al*. [17].

### Geomolecular dating with MCMCtree

All timetrees based on empirical and simulated data were inferred with MCMCtree, within PAML [28]. Both the independent rates and autocorrelated rates models were employed for the 122-taxon and 128-taxon empirical datasets, with control-file priors and run parameters replicating Springer *et al.* [10]. This includes unit time (100 Ma), the rate prior parameters, rgene_gamma shape (1) and scale (5.41), and the rate drift prior parameters, sigma_gamma shape (1) and scale (4.207). Analyses were run for 200,000 generations, sampled every 50^th^ generation, and a burnin of 10,000 generations was discarded. The 57-taxon alignments were similarly run in MCMCtree, although matching the original priors used by Liu *et al*. [17], including root age (4.16 - 4.254 Ma) and rgene_gamma (2, 40).

Rate distributions for the simulated datasets were based on the empirical data of Meredith *et al.* [2], with rate estimates taken from Phillips ([11], Fig. 5; also see “Results & Discussion”). In the case of the “ancestral rates” simulations, rates were modelled only from branches representing mammals <30kg adult body mass and <40 years maximum longevity. Given the rate estimate for these data of 0.1469 subs per 100 Ma, the rgene_gamma scale parameter was adjusted to 6.81 (=1/0.1469). The “mixed rates” analyses include four 33 Ma branches with 1/5 the ancestral rate, and as such the rgene_gamma scale parameter was adjusted to 7.51. The root age prior for all analyses of simulated data was 200 Ma (with sigma_gamma scale 2.0), with the root being symmetrically calibrated with soft 2.5% prior minimum and maximum bounds from 180-220 Ma.

### Eutherian mammal diversification in the fossil record

Direct reading of the eutherian fossil record implies an extraordinary diversification immediately following the 66 Ma KPg event [7, 8, 11]. However, Springer *et al.* [10] advocate other diversification spikes well before the KPg, during the Turonian (93.9-89.3 Ma) and Campanian (83.6-72.1 Ma). They also suggest another diversification spike after the KPg, during the Ypresian (56.0-47.8 Ma). However, it is important to consider fossil sampling. A stage with a short duration and poor sampling will artefactually appear to have few new species appearances, while the same actual diversification rate will result in many more new species appearances for a longer, and better sampled stage, especially if it follows a stage in which new appearances were masked by poor sampling.

We obtained fossil species richness and new appearance counts from The Paleobiology Database (accessed 29 March 2017). To help even out sampling potential we start with the critical (and well-sampled) Campanian, Maastrichtian and Paleocene, and then provide further time bins as individual or combined stages that sample at least 80 mammal species. Mammals overall provide a better indicator of sampling potential than eutherians, which are expected to have very low species richness close to their origin. Our strategy resulted in relatively even bin durations (average 8.78 Ma, s.d. 2.17 Ma, see Fig. 1, Supplemental file 5), except for the oldest bin, Albian-Cenomanian (19.1 Ma duration), which is outside the range of molecular and morphological predictions for the diversification spike. A second important factor that Springer *et al.* [10] did not consider for either fossil or molecular diversification analysis is the standing diversity base from which new fossil appearances derive, or from which new molecular lineages diverge, as is standard in lineage through time analysis (see [16, 61]).

Our metric for eutherian diversification is the number of new eutherian species appearances for the time bin, divided by both the duration of the time bin and the standing diversity of eutherians in the previous time bin. Fossil sampling potential is still unlikely to be constant across all of the time bins. Therefore, to integrate out much of the sampling disparity we also show new eutherian appearances in each bin as a proportion of new appearances among all mammals (indicated by colour scaling in Fig.1, also see Supplemental file 5).

## Supplemental files

**Supplemental file 1:** Addressing claims of “zombie” lineages on Phillips’ (2016) timetree.

**Table S1** and **Figure S1**. (pdf 661 kb)

**Supplemental file 2:** Fossil calibration schemes. **Table S2**, and **Figure S2**. (pdf 694 kb)

**Supplemental file 3:** MCMCtree timetrees. **Table S3**. (pdf 545 kb)

**Supplemental file 4:** GC3 conservation and estimating maximum longevity. **Figure S3**. (pdf 251 kb)

**Supplemental file 5: Table S4**. Fossil record species richness for Eutheria and Mammalia from Albian through to Lutetian. (pdf 91 kb)

## Acknowledgements

We are grateful to David Penny and Peter Waddell for valuable discussions on mammalian diversification and to Robin Beck for helpful comments on the manuscript. Thomas Halliday kindly provided dated morphological trees. This research was funded by an Australian Research Council Discovery grant (DP150104659 to MJP).

